# Core Defense Hotspots within *Pseudomonas aeruginosa* are a consistent and rich source of anti-phage defense systems

**DOI:** 10.1101/2022.11.11.516204

**Authors:** Matthew C. Johnson, Eric Laderman, Erin Huiting, Charles Zhang, Alan Davidson, Joseph Bondy-Denomy

## Abstract

Bacteria use a diverse arsenal of anti-phage immune systems, including CRISPR-Cas and restriction enzymes. Identifying the full defense repertoire of a given species is still challenging, however. Here, we developed a computational tool to broadly identify anti-phage systems, which was applied to >180,000 genomes available on NCBI, revealing *Pseudomonas aeruginosa* to possess the most diverse anti-phage arsenal of any species with >200 sequenced genomes. Using network analysis to identify the common neighbors of anti-phage systems, we surprisingly identified two highly conserved core defense hotspot loci (cDHS1 and cDHS2). Across more than 1,000 *P. aeruginosa* strains, cDHS1 is up to 224 kb (mean: 34 kb) with varied arrangements of at least 31 immune systems while cDHS2 has 24 distinct systems (mean: 15.4 kb). cDHS1/2 are present in most *P. aeruginosa* isolates, in contrast to highly variable mobile DHSs. Most cDHS genes are of unknown function potentially representing new anti-phage systems, which we validated by identifying a novel anti-phage system (Shango) commonly encoded in cDHS1. Identification of core gene markers that flank immune islands could be a simple approach for immune system discovery and may represent popular landing spots for diverse MGEs carrying anti-phage systems.

## INTRODUCTION

Bacteria are subjected to a diverse community of bacteriophages (phages) and rely on various anti-phage immune systems to block phage replication. Understanding the immune response to phage will be central to phage therapy success and likely new biological mechanisms await discovery. Anti-phage defense systems can be found adjacent to other defense systems within the same genomic locus in so-called “defense islands” (1). Defense islands are a rich source of novel anti-phage systems (2,3) with some showing homology to human and plant immune systems (4). For example, a bacterial version of Gasdermin, an innate mammalian immune system that forms large pores in the cellular membrane, was recently discovered by searching for gene families enriched in defense islands (5). CBASS is another remarkable example of an anti–phage immune system found in defense islands, which is homologous to cGAS-STING innate immunity (6).

Defense islands are often found peppered across the genome with no conserved positioning, due to being frequently encoded by mobile genetic elements (MGEs). Numerous reports have documented the phenomenon of the dissemination of mobile anti-phage immunity as a driver of phage resistance in the wild. For example, isolates of *Vibrio lentus* and *Vibrio cholerae* have MGE-disseminated islands that contain multiple anti-phage systems in hotspots to protect from phage (7, 8). Additionally, *E. coli* P2-like and P4-like prophages encode several anti-phage systems at a single hotspot that antagonize competing phage (9). These discoveries enable the facile identification of immune systems in *these* MGEs and in genomes where they are found but the complete identification of immune systems in a genome or species remains an important bioinformatic and experimental challenge.

Here, we sought to comprehensively identify and annotate the defense systems of *Pseudomonas aeruginosa*, a generalist microbe, opportunistic human pathogen, and a model organism for phage-host interactions and CRISPR-Cas biology. These efforts lead to the unexpected observations of two conserved loci in most *P. aeruginosa* genomes that serve as hotspots for immune systems, which we call core defense hotspots (cDHS). We show that cDHS1 contains at least 31 different immune systems across 1,672 isolates, with many genes of unknown function likely involved in phage defense. One operon that is commonly present in cDHS1 (a new system we call “Shango”, containing a TerB and helicase domain) inhibited replication of several phage families. In contrast to defense islands encoded on variable MGEs, this core DHS is encoded in the same region of the host chromosome in all isolates and appears to have been assembled by numerous diverse MGEs. We also find another conserved hotspot with extensive immune diversification called cDHS2. These regions are likely rich in new anti-phage systems and do not require any specific existing immune system or single MGE family to find them. Core defense hotspots might be a common phenomenon in bacteria and network analyses like ours provide a road map to their discovery.

## MATERIAL AND METHODS

### Annotating *P. aeruginosa* immune systems with ISLAND pipeline

ISLAND uses Hidden Markov Models (HMMs) and BLAST searching to annotate anti-phage systems in large bacterial genome databases. HMMs were created by taking a set of seed protein sequences, removing all redundant sequences, and clustering them using MMseqs2 *easy-cluster* at high sensitivity (*-s 7*.*5*). A multiple sequence alignment from each cluster with more than five members was then created using Clustal Omega, and HMM models were built using hmmer (hmmer.org). HMMs for CRISPR-Cas systems were obtained from CRISPRCasFinder and matches to these HMMs were searched for using macsyfinder (10, 11). Representative sequences for immune systems from BREX, DISARM, Gabija, Wadjet, Septu, Shedu, Thoeris, Druantia, Lamassu, Hachiman, Kiwa, Zorya, CBASS, Retrons, *etc*., were obtained from literature (2, 3, 12, 13). Representative sequences for Type I, II, III, and IV RM systems were obtained from REBASE’s gold standard collection (14). Type II and IV RM systems were searched for by BLAST as effective HMMs cannot be built for these systems (15, 16). The sequences for each gene in an experimentally verified system consisting of at least two genes were PSI-BLASTed against all genomes available on NCBI in GPFF format (12/03/2020) with four iterations and an inclusion e-value threshold of 0.005. Systems were identified by searching for hits (e-value < 1e-10) of each gene that were directly adjacent to a hit corresponding to its neighbor in the system it is from and part of a locus where all genes from a given system were accounted for. HMMs for all genes from each system were created from the PSI-BLAST hits that were determined to be part of a bona fide system using the method described above.

Immune systems were found by searching for all genes comprising that immune system using hmmsearch or BLAST (only if an HMM could not be built for that gene). Searches were performed against databases in gembase format. The database was first clustered with mmseqs2 easy-linclust with an identity cutoff of 0.9 (17). Only representative sequences for a cluster were included in the temporary database. All genes for systems of interest were searched for in the temporary database using hmmsearch or BLASTP as appropriate. If a protein in the temporary database was found to be a hit for a gene in a system of interest, all other proteins in the original database that belong to the same cluster as that protein were considered hits for that gene. Once hits for all genes of interest were identified, systems were identified. A system was required to have all essential genes belonging to that system and enough accessory genes belonging to that system to reach the minimum number of required genes for that system found within a system-specific number of genes of each other (Figure 1).

**Figure 1:**
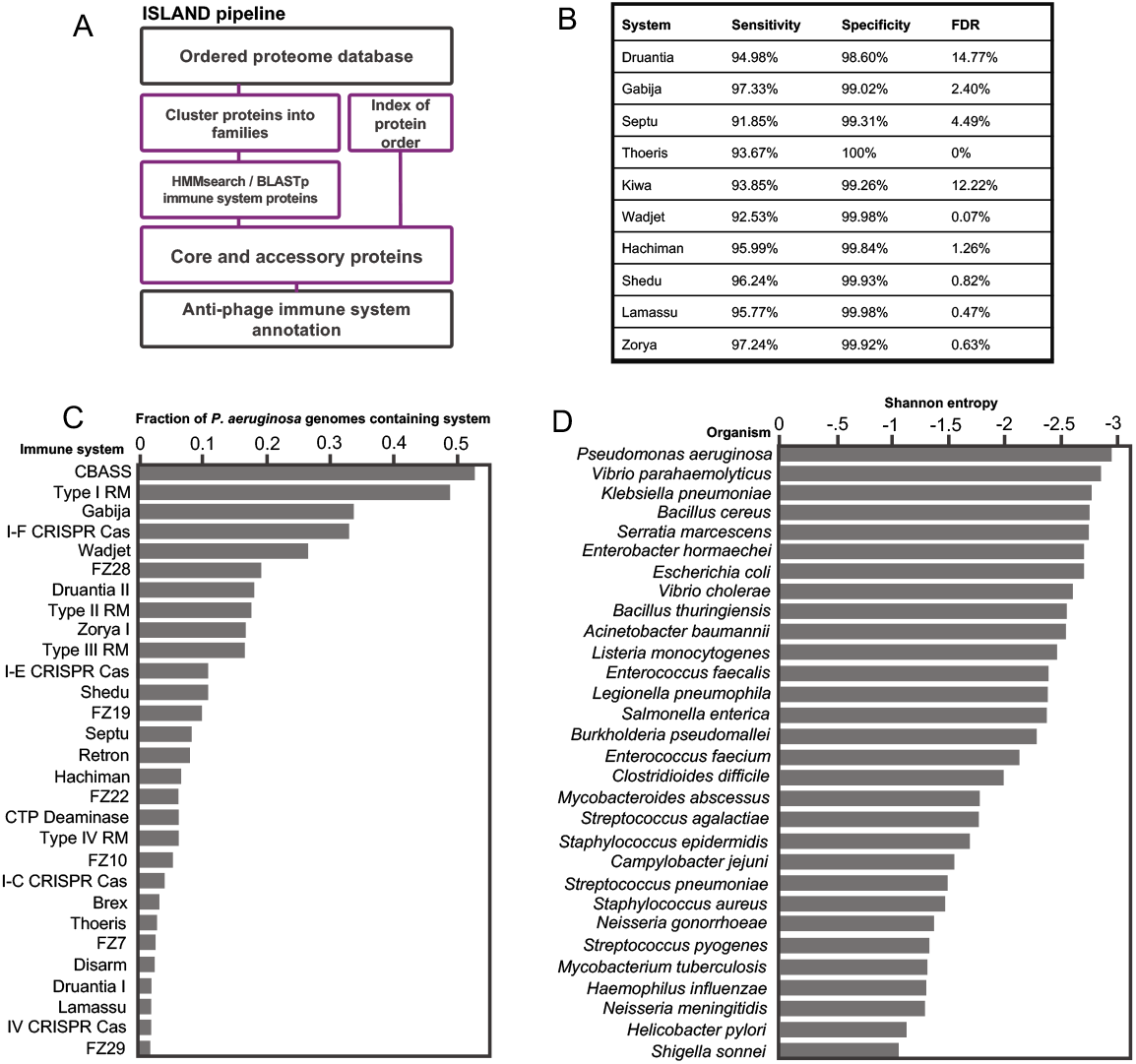
ISLAND workflow system distribution/diversity. (A) The ISLAND workflow requires building HMMs for genes in anti-phage systems, searching a pre-clustered protein database, and applying pre-defined logic. Models for genes in a system of interest are created by obtaining protein sequences from the literature corresponding to this system, clustering them, and then building HMMs from the clusters with at least three genes. Then systems are searched for in an ordered protein database (gembase format). (B) The sensitivity/specificity and false discovery rate of ISLAND of the Doron et al., 2018 systems (C) The fraction of *P. aeruginosa* genomes that contain each of the antiphage systems shown. Only anti-phage systems present in at least one percent of *P. aeruginosa* genomes available on NCBI were shown above. (D) Shannon entropy as an indicator of immune system diversity across bacterial species with at least 500 genomes in NCBI.

### Network analysis

The protein sequencing making up the genomic loci 5 kb upstream and downstream of the immune systems annotated by ISLAND were retrieved. For each set of proteins comprising the locus, a subset list of all combinations between the identified system and the proteins in the locus was created. The pool of proteins in all defense islands were clustered using MMseqs2 linclust tool with default parameters. This produced protein clusters or families. These protein families were mapped back to the combination subset lists and embedded in a directed network where system nodes are connected to protein family neighbor nodes. Cytoscape 3.9.1 (18) was used to visualize the resulting network.

### cDHS1 and cDHS2 region analysis

To find cDHS1, two conserved genes were chosen for their conservation *Pseudomonas*. The protein sequences of the MerR transcriptional regulator and the DUF4011 protein of PA14 were used as markers of the left and right boundary respectively. MMseqs2 search function was used to find orthologs in our database downloaded from www.pseudomonas.com. Contigs were filtered for having both markers. Data on these contigs and their hotspot can be found in supplementary table 1. cDHS2 locus was retrieved by using the tmRNA site and the VOC, LysR, RidA like proteins as markers.

### Phylogenetics of cDHS1

To construct a phylogeny of the *Pae* isolates, the redundant nucleotide sequences of cDHS1 were removed using MMseqs2 and the entire genome for the remaining set were retrieved. Gubbins (https://github.com/nickjcroucher/gubbins) was used to construct a RAxML maximum likelihood tree. Gubbins was chosen because it accounts for extensive horizontal transfer by removing hypervariable regions before alignments. iTOL (https://itol.embl.de) was used to apply features of the cDHS1 to the tree To identify long range synteny, tree branch distances were calculated between every pairwise isolate using the Phylo package from BioPython. The cDHS1 nucleotide MASH-distance was calculated between every pairwise isolate using MASH (https://github.com/marbl/Mash). Long range synteny was defined as isolated having an evolutionary tree distance above 0.35 and an cDHS1 MASH-distance above 0.9. Occurrences of rapid diversification were defined as isolates having a tree distance below 0.05 and a cDHS1 MASH-distance above 0.38.

### Domain analysis

To better understand the molecular function of the genes encoded at these hotspots, domain analysis was performed on all protein-coding genes within the boundary genes. Pfam domain models were obtained from Conserved Domain Database (May 3rd, 2022) and searched against the pool of proteins within cDHSs that were not annotated by defense-finder as belonging to an immune system. Rpsblast (2.11.0+) using an e-value cutoff of 0.01 was used to search for the pfam models.

### Mobilome analysis

MobileOG-db was used to quantify mobile elements in cDHSs (19). Sites were separated into three categories (integrative, prophage or conjugative) if they matched ‘integration’, ‘phage’, ‘conjugation’ in their major category annotation.

### Strain and plasmid construction

cDHS1 of PA14 contains Shedu (PA14_RS11720) and other defense-like proteins, including a large multi-domain helicase (PA14_RS11730), a three-gene TerB-like protein (PA14_RS11735), ATPase (PA14_RS11740), and DEAD/DEAH box helicase (PA14_RS11745) system, referred to as Shango. The three gene system including its native promoter with ihfA and MerR genes found in the flanking regions of cDHS1 (Figure 3A) was cloned in pHERD30T digested with PstI-HF and SacI-HF enzymes for 1 hour at 37C. PCR was used to amplify the system from PA14 and subsequently assembled in digested pHERD30T using Gibson assembly (pMJ31). Phage sensitive host PAO1 was chosen because it is sensitive to several phages. pMJ31 was electroporated into PAO1 and recovered with SOC for 1 hour and plated on gentamicin 50ug/ul.

### Phage assays

Phages were 10-fold serial diluted 8 times in SM phage buffer. 2ul of the phage dilutions were plated on PAO1 either containing pHERD30T empty vector or pMJ31 (pHERD30T-Shango) that were 150ul diluted into 3ml of 0.7% top agar. Plates were left to try under a flame for 10 minutes and incubated upside down, to prevent condensation, at 37C overnight. Plate images were taken with a Biorad Gel Doc EZ imager. EOP was calculated as a ratio of the number of plaque forming units (PFUs) that formed the strain expressing Shango divided by the number of PFUs that formed on the strain with an empty vector. Liquid culture phage infections were prepared with strains grown overnight and diluted 100x in LB supplemented with 10mM MgSO4, gentamicin, and 0.3% arabinose. 140ul of bacterial culture was infected with 10ul of serially diluted phage in a 96-well plate. Growth was monitored over 35 hours using a Synergy H1 microplate reader (BioTek) shaking at 37C.

## RESULTS

### Immune system detection with ISLAND is specific and sensitive

To enable the identification of anti-phage bacterial immune systems, we developed a bioinformatic pipeline called Immune System Loci Annotation ‘n’ Database (ISLAND). ISLAND uses a combination of hidden Markov models (HMMs) and BLASTp to annotate immune systems in large bacterial genome databases (Figure 1A). To test the annotation accuracy ISLAND, we measured our pipeline’s ability to identify the systems included in Doron et al., 2018. We included only the genomes that were found to have at least one system in Doron et al. 2018 and were available from NCBI (14,296 genomes). Our models had very high sensitivity and specificity, and generally low false discovery rates (Figure 1B).

We then applied ISLAND to all genomes available on NCBI as of 12/03/2020 (184,541 genomes). Across all species we found that the most common systems were RM systems with more than 70% of all genomes surveyed containing at least one RM system. The three next most common systems were CRISPR-Cas, CBASS, and Gabija, which were found in 35%, 16%, and 10% of genomes, respectively. Our results on overall immune system abundance are qualitatively similar to those recently presented in Tesson et al 2022 (Figure S1). We found fewer anti-phage systems per genome than Tesson et al., however, as our database contained models for fewer systems.

### *Pseudomonas aeruginosa* as a model for immune system discovery and gene organization

For each species with over 500 genomes deposited on NCBI we calculated the Shannon entropy of defense as the sum of p_sys_*ln(p_sys_) for each system present in the species. p_sys_ is the proportion of all defense systems present in a species that are a given system. We found that *P. aeruginosa* has the most negative Shannon Entropy of defense, indicating that its complement of immune systems is the most diverse of any of the bacterial species analyzed. In *P. aeruginosa*, CBASS and Type I RM are the most common anti-phage systems, present in 53% and 49% of genomes respectively. Gabija, Type I-F CRISPR-Cas, and Wadjet are the next most common systems, and all are present in more than 20% of *P. aeruginosa* genomes (Figure 1C). Furthermore, of the ten most common anti-phage systems in *P. aeruginosa* only four (Type I and II R-M, Type I-F and I-E CRISPR-Cas) had been known before 2018, suggesting that there are many antiphage systems waiting to be discovered. Other bacterial species with highly diverse immune arsenals include *Escherichia coli, Klebsiella pneumoniae*, and *Vibrio parahaemolyticus*. On the other hand, *Campylobacter jejuni, Heliobacter pylori, and Neisseria meningitidis* all have immune arsenals which are not nearly as diverse by Shannon entropy (Figure 1D). The immune system prevalence and diversity in *P. aeruginosa* coupled with genetic tractability and a diverse isolated phage population suggests that *P. aeruginosa* will be a good model organism for the characterization and discovery of new anti-phage immune mechanisms.

### Core defense hotspot 1 (cDHS1) in *P. aeruginosa*

After validating the accuracy to annotate immune systems with ISLAND, the pipeline was applied to the *Pseudomonas* database (www.pseudomonas.com) containing at least 4,640 *P. aeruginosa* genomes. Specifically, our goal was to understand the complement of anti-phage systems in *P. aeruginosa* and where in the genome they are encoded. The genes found within 10 kb surrounding a known immune system (i.e., putative defense islands or genes flanking a defense island) were retrieved from the output and encoded as a network (Figure 2). This resulted in a large, bipartite network with most nodes having a degree of one (gene neighbors only found nearby a single known immune system) and several nodes having a high degree (genes associated to several systems immune system). We considered that nodes with the highest degree might be candidate immune systems. This was not the case however, as we observed a clear cluster of gene nodes with functions related translation: Phenylalanine tRNA ligase subunit alpha (PheS), Phenylalanine tRNA ligase (PheT) subunit beta, and 50S ribosomal L20 (rpIT)) that were associated with nine different immune systems (Figure 2). *pheT* is essential in *P. aeruginosa* and broadly conserved (20). This region also contained a MerR family transcriptional regulator, an integration host factor gene, ihfA, and a proline tRNA.

**Figure 2:**
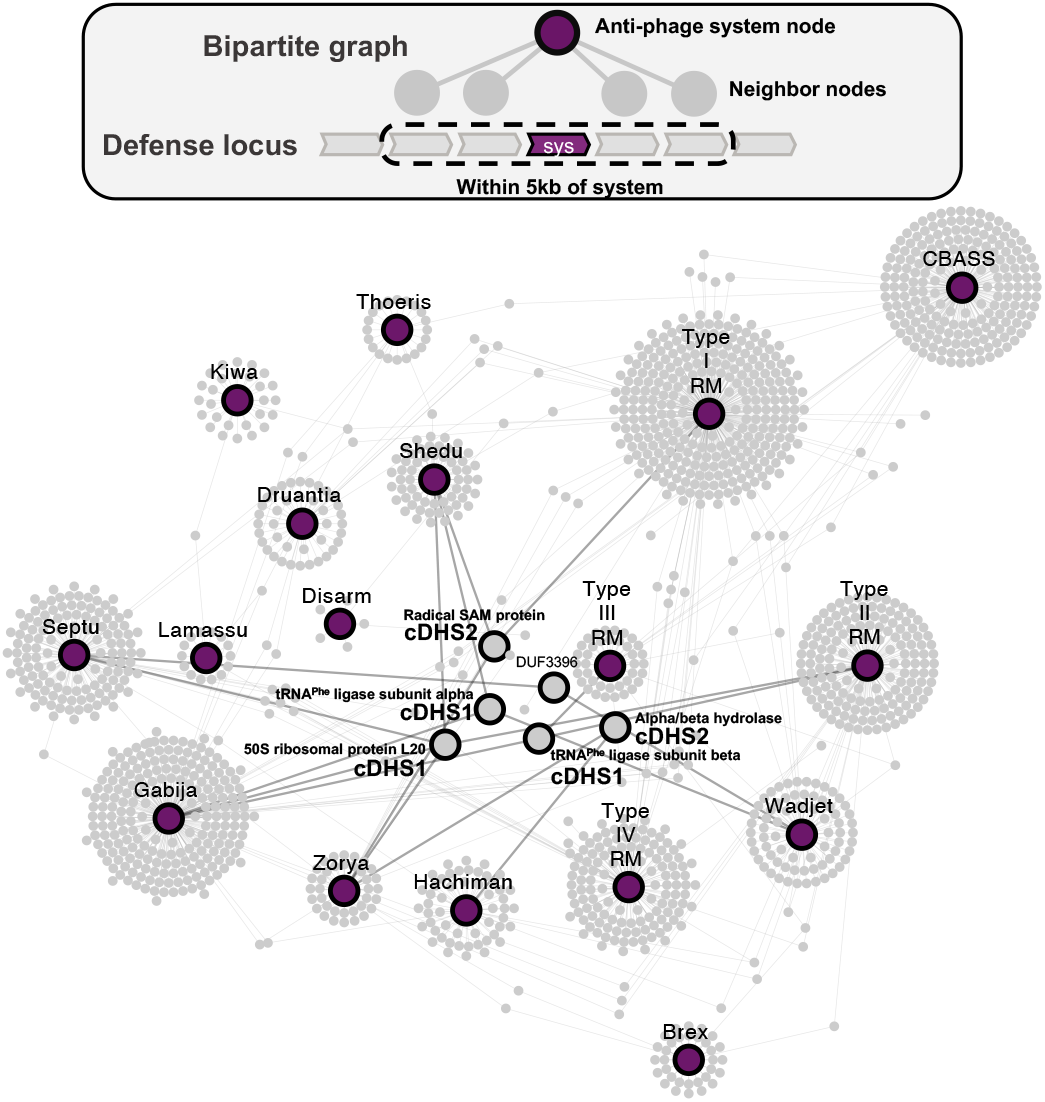
Defense island association network. Purple nodes represent anti-phage systems and gray nodes represent genomic neighbors of that system within 5 kb upstream and downstream. Most gray nodes are only represented nearby one system and are those shown acting as a “halo” around the purple nodes. Other nodes are present nearby multiple systems and are towards the center of the network. The large gray nodes with black borders are nodes of interest because they represent genes that are core/conserved and associated with many different immune systems.

These genes related to translation appeared as the left boundary to a locus that contained numerous immune systems (Figure 3A), with the right boundary containing a conserved small hypothetical gene, a large DUF4011 gene, and a radical SAM protein which was also a node of interest in our network (Figure 2). To get a sense for how conserved this locus is in *P. aeruginosa*, homologs of the MerR family protein and the DUF4011 were searched across our database. These markers were found in every *P. aeruginosa* isolate in our database. We therefore consider this region to be a defense hotspot (DHS) that is marked by consistent flanking core genes and not obviously a member of any one group of mobile genetic elements (MGEs), which we dub as core DHS1 (cDHS1). Genomes with both markers on the same contig were filtered and resulted in more than 1,700 total cDHS1 regions. cDHS1 ranged dramatically in size (1-224kb with a mean of 34k kb, Figure 3C). The region has a lower GC% content compared to the *P. aeruginosa* genome (58.9% vs. 65.3%) (Figure 3D).

**Figure 3:**
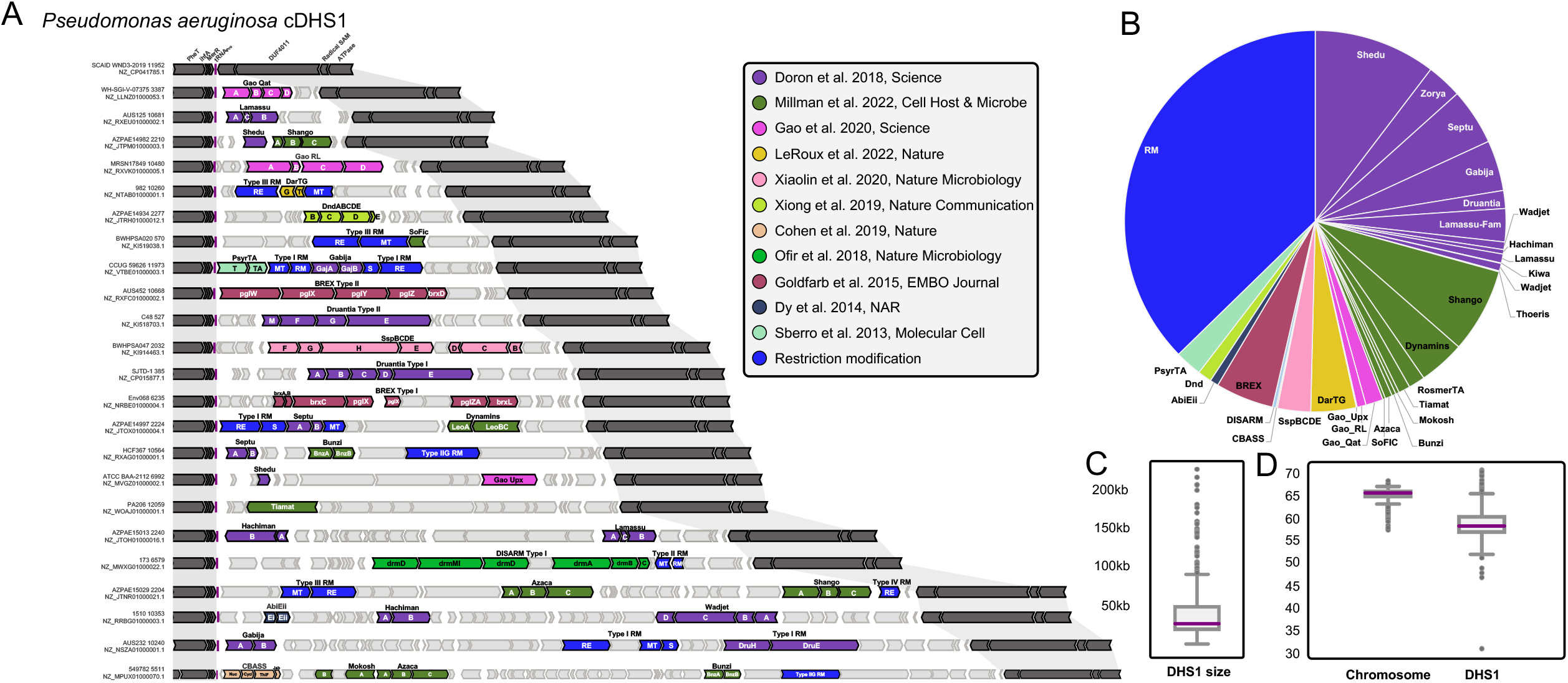
Core defense hotspot 1 (cDHS1). (A) Representative isolates with cDHS1 chosen to show the diversity in anti-phage systems. Dark gray genes are marker genes that define the boundaries of the hotspot. Light gray genes indicate non-defense genes and are neighbors to the defense system which are colored based on the publication they were discovered in. (B) Pie chart of the different immune systems found in cDHS1, colored by publication. (C) Size distribution of cDHS1 in kb. (D) GC% difference between the host chromosome and the defense hotspot.

DefenseFinder, a recently published anti-phage annotation tool (21) was used to annotate the immune content of this locus as it contains the most comprehensive set of immune system HMM profiles to date and is orthogonal to our ISLAND program used to initially identify this locus. DefenseFinder annotated 31 different immune systems at this site, a remarkable hotspot for anti-phage immunity. The ubiquitous nature of RM across prokaryotic genomes persists in cDHS1. Of the total number of defense systems in cDHS1, Shedu, Type I RM and Type III RM were the most common immune systems found, with over one third belonging to RM (Figure 3B). We find that another large percentage of immune systems come from publications that used computational analysis of defense islands (2, 3, 22) (Figure 3B). We find at least one immune system in 83.7% of cDHS1 loci (Supplemental table 1), clearly demonstrating an immune functionality for cDHS1. While this is a strong anti-phage hotspot, 87% of genes in the cDHS1 locus are not a part of a known immune system, suggesting this locus will serve as a rich source of novel discovery.

### Core defense hotspot 2 (cDHS2) in *P. aeruginosa*

Since the network revealed cDHS1 through high degree nodes, we asked if other high degree nodes would point to another hotspot. We noticed an alpha/beta hydrolase annotated node that pointed us to another defense rich locus in *P. aeruginosa*. We performed the same analysis for this locus as we did for cDHS1 and found it to exhibit features like cDHS1, we denote this site cDHS2 (Figure 4A). cDHS2 contains 24 different immune systems, less than that of cDHS1, but three systems were present in cDHS2 that were not in cDHS1 (Retron, PARIS, and dCTP deaminase). It showed clear hypervariability and diversification with a range in size between 0.3 kb to 168 kb (Figure 4B). It also has a different composition of immune systems. While restriction modification is the chief system at both cDHS1 and cDHS2, Druantia and Wadjet account for roughly 25% of the immune systems within cDHS2, while these systems are rare (<2%) in cDHS1. Conversely, systems Shango and BREX make up ∼12% of systems in cDHS1 but Shango is not found and only two occurrences of BREX (<1%) in cDHS2. Similar to cDHS1, this site is flanked by a non-coding RNA, in this case an *ssrA* transfer-messenger RNA.

**Figure 4:**
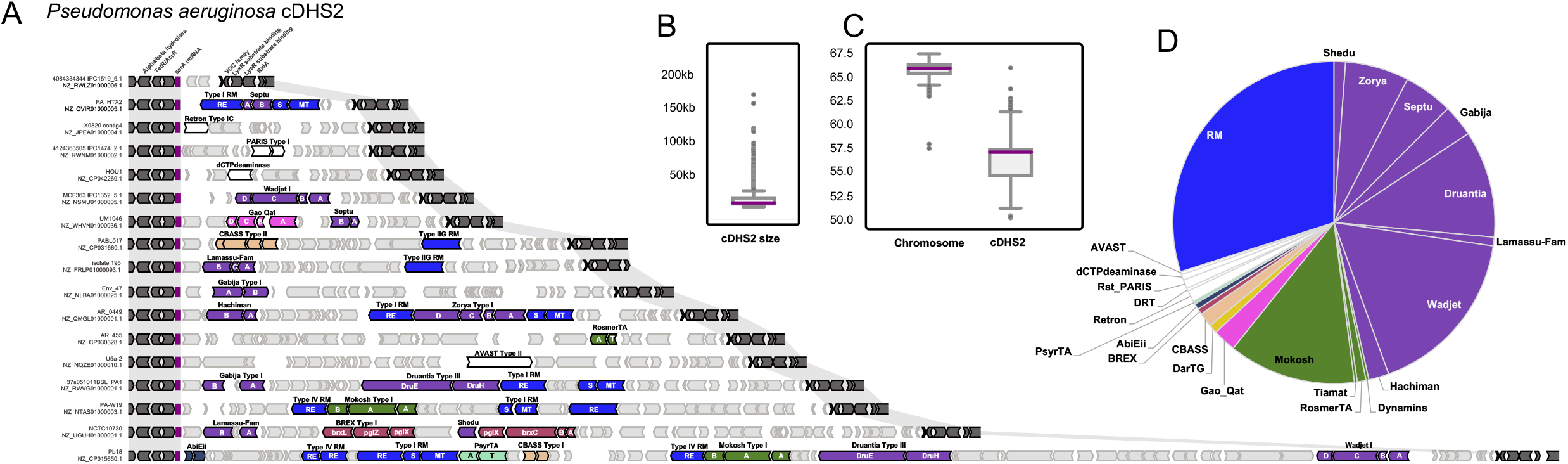
Core defense hotspot 2 (cDHS2): (A) Representative isolates with cDHS2 showing diversity in anti-phage systems (same color scheme as figure 3) (B) Size distribution of cDHS1 in kb. (C) GC% difference between the host chromosome and the defense hotspot. (D) Pie chart of the different immune systems found in cDHS2 (same color scheme as figure 3).

### Domain analysis

To understand what other genes might be in these hotspots and to search for potential mobilizing genes, we performed Pfam domain analysis on all the genes not annotated by ISLAND or DefenseFinder. While most genes are hypothetical and have no known domains, we find many domains related to anti-phage defense (toxins, helicases, ATPases, etc.,) (Figure 5A). A *symE* toxin is commonly found among cDHS1 sites. *symE* is small toxin belonging to the Type I toxin-antitoxin systems characterized in *E. coli*, the cognate antitoxin is an antisense RNA that binds to the 5’ untranslated region (UTR) of *symE* (23, 24). This TA module could be stabilizing the locus as an addiction module, ensuring the acquisition of subsequent immune systems in cDHS1, or perhaps it has anti-phage activity. Domain analysis of cDHS2 paints a different story. Most genes (87%) contained no known pfam domains, but of those that had domains, many of them appear to belong to a membrane transport system which might belong to conjugative systems (Figure 5B).

**Figure 5:**
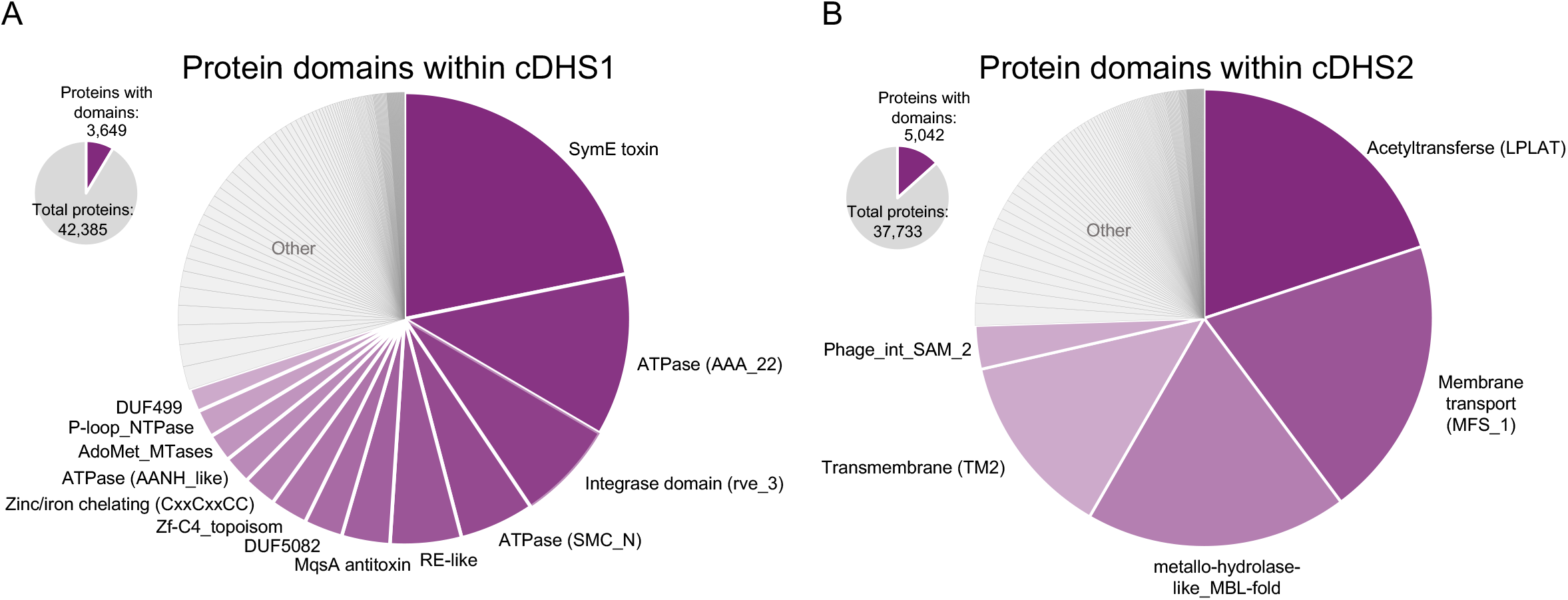
Domain analysis of cDHS1 and cDHS2: Pie chart summarizing the non-defense genes in each hotspot. Note that most genes do not match a pfam domain. Of the genes with domains, the most common are indicated in purple for cDHS1 (A) and cDHS2 (B).

### Shango is an anti-phage system

We noticed a three gene system common in cDHS1, including model strain PA14. This system contains a TerB-like protein, a helicase, and an ATPase. We cloned this system on a plasmid and expressed it in strain PAO1 (a strain with a small cDHS1) and performed phage plaque assays with several phages from different morphotypes (Podoviridae, Siphoviridae, and Myoviridae) in liquid and solid media (Figure 6). We see modest (i.e., 10-100-fold EOP change) anti-phage activity against many phages on solid media, (Figure 6B). For several phages, we noticed a reduction in plaque size and/or clarity, suggesting decreased phage fitness. Notably, this was not observed for all phages titrated on lawns expressing Shango (Figure S2). Generally, we see strong protection of cell cultures from phage-induced lysis over a broad range of phage concentrations during liquid growth (Figure 6C, S3). We call this anti-phage system, Shango. Shango was recently identified in immune islands in *E. coli* and modestly inhibited the replication of two different phages by ∼10-fold (22).

**Figure 6:**
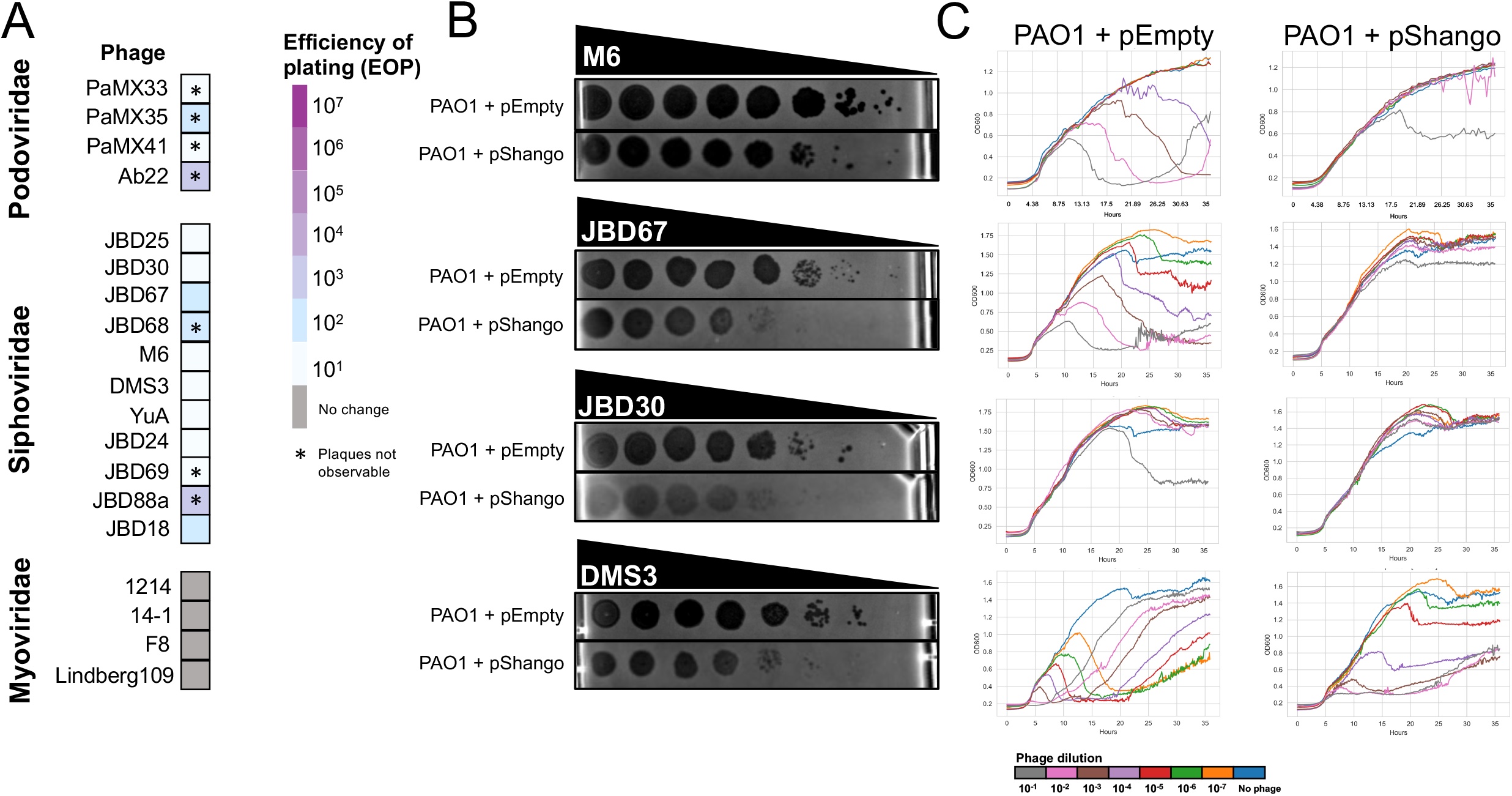
Shango phage assays: (A) A heatmap of efficiency of plating (EOP) of several different *Pseudomonas* phage on a strain. EOP is calculated as the ratio of the number of plaques between the PAO1+pEmpty (empty vector) strain and the PAO1+pShango expressing strain. Phage are grouped by their morphology type (*Myoviridae, Siphoviridae*, and *Podoviridae*). (B) Ten-fold dilutions of phage lysates (spotted left to right) on a lawn of PAO1+pEmpty or PAO1+pShango. (C) The same strains and phages were used in liquid infection, where OD600 is plotted to quantify culture lysis due to phage replication. Phage dilutions are labeled at the bottom.

### Core DHS1 undergoes rapid immune adaptation

To understand the cDHS1 immune diversification across *P. aeruginosa* isolates, a whole genome phylogeny was constructed of all isolates containing an intact cDHS1 (Figure 7A). Evolutionary discordance between the chromosome and cDHS1 is expected if these sites are horizontally acquired (Figure 7B). Indeed, we see this discordance amongst cDHS1 sites in the tree, where isolates in different clades of the tree have high cDHS1 similarity at the nucleotide level with one or two systems differentiating the two (Figure 7C). Moreover, we find examples of acquisition of new immune systems in the middle of cDHS1. While isolates that are very closely related tend to have similar cDHS1 sites, occasionally there is rapid diversification of the hotspot (Figure 7D) that can gain several kilobases. Conversely, we see synteny between cDHS1 sites across distantly separated isolates. These data confirm what is expected given the diversity of systems at this single locus, that this region is likely mobilized and diversified through horizontal gene transfer. One potential route would be phage transduction and recombination, since the regions flanking cDHS1 are highly conserved in all isolates.

**Figure 7:**
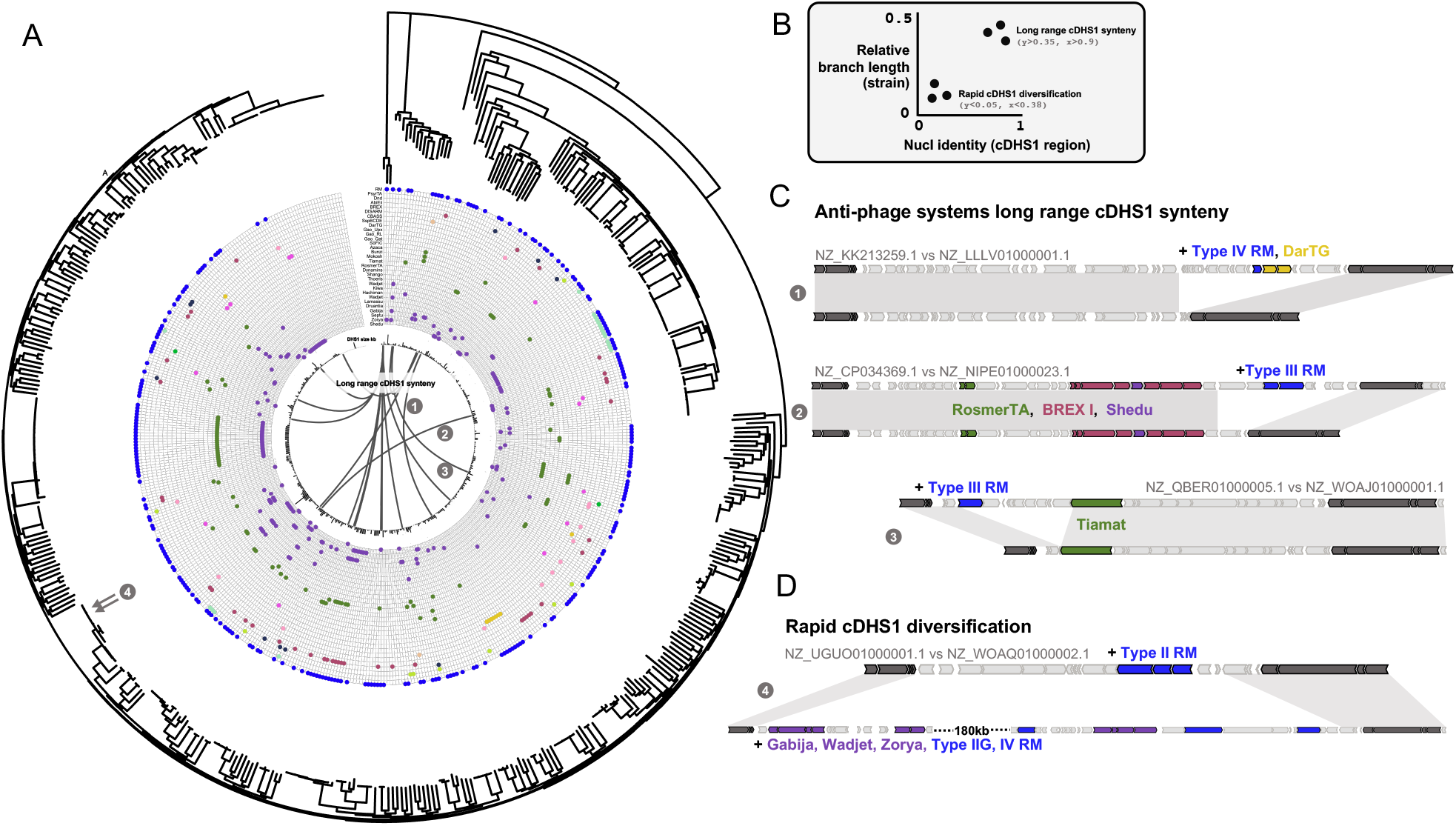
Phylogeny and synteny of cDHS1: (A) A RAxML tree of the isolates encoding cDHS1 with the systems within the hotspot indicated by a colored dot (same color scheme as figure 3). cDHS1 size is represented by the black bars in the center of the tree. Lines between pairs of isolates indicate long range synteny which is exemplified on the right side. (B) Example plot showing how long range synteny and rapid diversification is determined. Pairs of isolates with high branch length between their points on the tree (distant isolates) and high nucleotide identity of cDHS1 indicate long range synteny. Pairs with low branch lengths (close isolates) but high cDHS1 identity indicate rapid diversification. (C) Anti-phage systems long range synteny with the same color scheme as figure 3. Light gray ribbons represent high nucleotide identity. (D) Rapid cDHS1 diversification where two isolates with very close proximity on the tree have extremely different cDHS1 compositions.

### Mobilome of cDHS

We next assessed the mobilome within genes between the cDHS flanking regions and found no single MGE type amongst the sites, rather a combination of transposons, conjugative elements, and prophages were identified. 29% of cDHS1 and 47% of cDHS2 sites had no detectable mobile gene (Figure 8A, B). Transposases were the most found mobilizing gene in both cDHSs, most of them belonging to the IS3 family (Supplemental table 1). Most transposases were found at the boundaries of the hotspots (Figure 8C). Prophages (full or partial) were found in a small percentage of cDHS sites as well as genes of viral origin, like the YgaJ viral recombinase. We also find examples of type IV conjugation systems and plasmid partitioning proteins in a small fraction of cDHS sites. The ability of these sites to accept a wide range of MGEs could be why cDHS are almost always occupied. Notably, the flanking “core” region of both cDHS1 and 2 possess non-coding RNA genes, which are popular landing sites for MGEs (25-28). cDHS sites therefore apparently act as a landing pad for mobilization events stemming from diverse MGEs encoding anti-phage immune systems.

**Figure 8:**
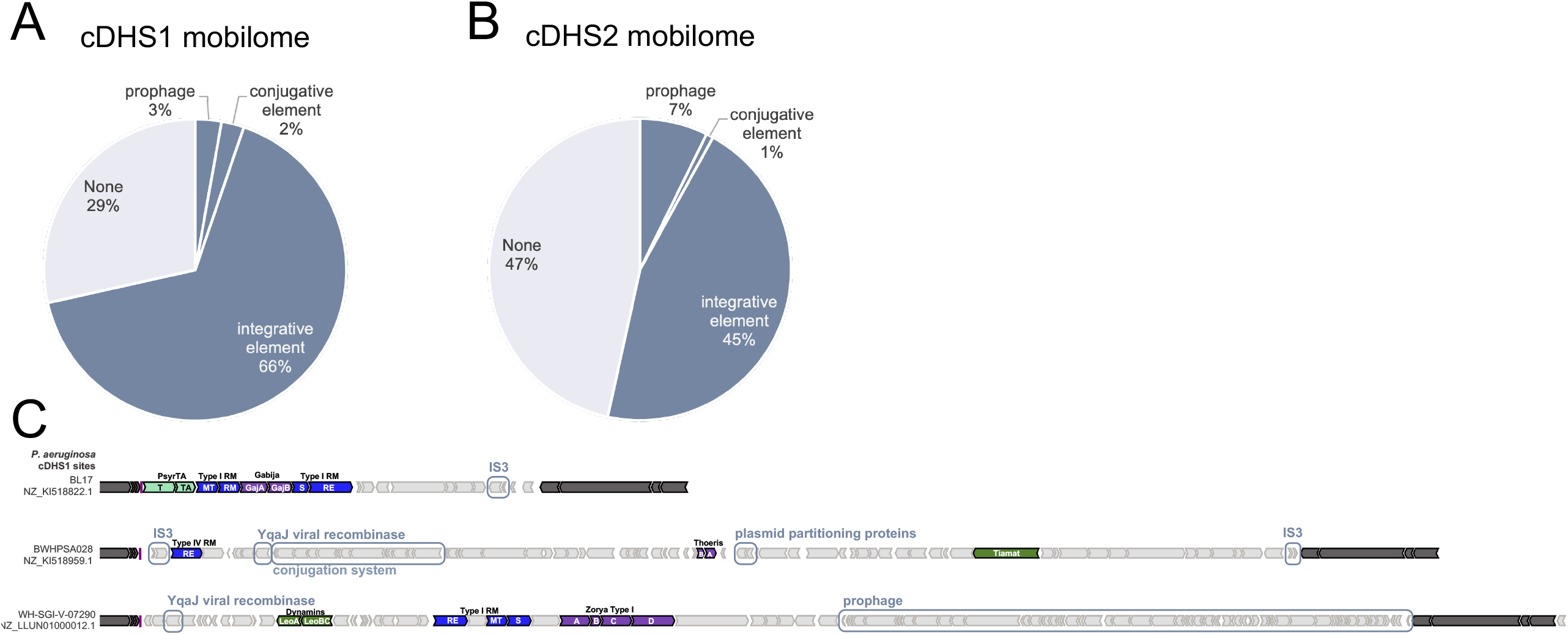
Mobilome of cDHS: Pie chart of the mobile element types that make up cDHS1 (A) and cDHS2 (B). (C) Orf maps of cDHS1 that carry mobile genes (boxed). Defense systems are colored based on figure 3.

## DISCUSSION

The current explosion anti-phage immune systems discovered in bacterial genomes has prompted the analysis their genomic positioning. Fascinating examples of MGE transferred hotspots of anti-phage immunity have been documented that control ecological outcomes during phage-host interactions in *Vibrio* and *E. coli*, for example (7-9, 29-31). In contrast to these phage defense MGEs, however, here we show that core defense hotspots (cDHS) with conserved genomic markers flanking immunity regions can be predictive of an immune locus. While immunity-carrying MGEs may integrate in some genomes but not others, cDHS regions by definition are essentially always present in all isolates of a species. This differentiation aligns with the reports of two types of flexible islands in clonal prokaryotes, replacement and additive islands. Replacement islands, like mDHS, are made up of short defense cassettes that share no sequence homology to each other. Additive islands, like cDHS, result from multiple discrete integration and are often associated with tRNA’s (32). An outstanding question not addressed here is whether cDHS regions are common in other bacterial clades, which will be addressed in future work.

In contrast, cDHS regions in *Pseudomonas aeruginosa* we sporadically identified mobilizing genes (e.g., transposases), the regions have a large size variance across the isolates, and we observe distant isolates carrying almost identical DNA between the cDHS flanking regions with the gain/loss of an anti-phage MGE (Figure 7C). In contrast, the elements present at 41 mDHS sites described in *E. coli)* typically belong to a single mobile genetic element family and their presence is hypervariable (i.e., each locus is occupied in 8% of isolates on average) (31). These are signatures of a landing spot for a specific MGE. A plausible three-step model could be at play where (i) cDHS contains a target site that acts to seed the initial insertion element that could (ii) carry additional target sites which facilitate subsequent insertion events and then (iii) dissemination the entire hotspot to other isolates. If the founding MGE that recognizes the target site bears an anti-phage system, it may stage the region with a bias to maintaining additional defense systems as is the case for defense islands (1). If the target site is disrupted after the initial integration event, only MGEs with target sites near or within defense systems would be compatible with the region thereafter. Indeed, we observe defense systems like Gabija, Septu, DarTG “inside” RM systems (Figure 3A). Finally, the entire region has potential to laterally transfer due to the conserved flanking regions. Transfer could be facilitated through generalized phage transduction and homologous recombination, as this would not require additional MGE-derived scars in the genome. How these enigmatic core defense hotspots elements are built, and whether they can transfer in this manner will be important areas for future work.

The identification of cDHS greatly simplifies the identification of known and new immune systems in *P. aeruginosa*. Notably, we show the utility of immune discovery via cDHS regions by identifying Shango from the cDHS1 of PA14 which is an anti-phage system with a TerB-like domain. It was previously hypothesized that the widespread nature of tellurite resistance (Ter) domains and its association to several anti-phage-like domains could be indicative of anti-phage mechanisms (33). Core defense hotspots identified here are present in nearly all strains of *P. aeruginosa* and >80% of the time have at least one known immune system and likely many others that await identification (i.e., >85% of genes present in cDHS1/2 are of unknown function). The systems published by Doron et al., in 2018 account for almost half of the systems found at cDHS2. During the preparation of this manuscript, another report with 21 new immune systems was published, including Shango (22). These systems account for roughly a quarter of the known immune systems in cDHS1 and cDHS2. The collection of immune systems in a discrete number of genomic loci also generates the possibility that expression is coordinated. Endogenous repressive pathways could be co-opted experimentally or as a phage therapy aid, to enhance phage infection in clinical pathogens (34)

The immune repertoire in *P. aeruginosa* is incredibly rich and we postulate that this organism will be a source of immune system discovery in the future. *P. aeruginosa* is additionally well-suited for studying the endogenous mechanisms, (alternative) functions, and regulatory strategies behind anti-phage immunity. The generalist lifestyle, broad ecological niches, and phage diversity likely necessitates an active pan-immune repertoire.

## DATA AVAILABILITY

ISLAND hidden Markov model can be found at DOI:10.5281/zenodo.7254690 (https://doi.org/10.5281/zenodo.7254690)

## SUPPLEMENTARY DATA

Supplementary Data are available at bioRxiv.

## ACKNOWLEDGEMENT

We would like to thank Sam Sternberg for his insightful comments on the manuscript. We also want to thank Sukrit Silas for early discussions regarding the design of ISLAND and Bondy-Denomy lab members for support and input.

## FUNDING

This work in the Bondy-Denomy lab was supported by the National Institutes of Health (NIH) (R01GM127489, R01AI171041, R21AI168811) and the Robert J. Kleberg, Jr. and Helen C. Kleberg Foundation.

## CONFLICT OF INTEREST

J.B.D. is a scientific advisory board member of SNIPR Biome, Excision Biotherapeutics, and LeapFrog Bio, and a scientific advisory board member and co-founder of Acrigen Biosciences. The Bondy-Denomy lab receives research support from Felix Biotechnology.

---

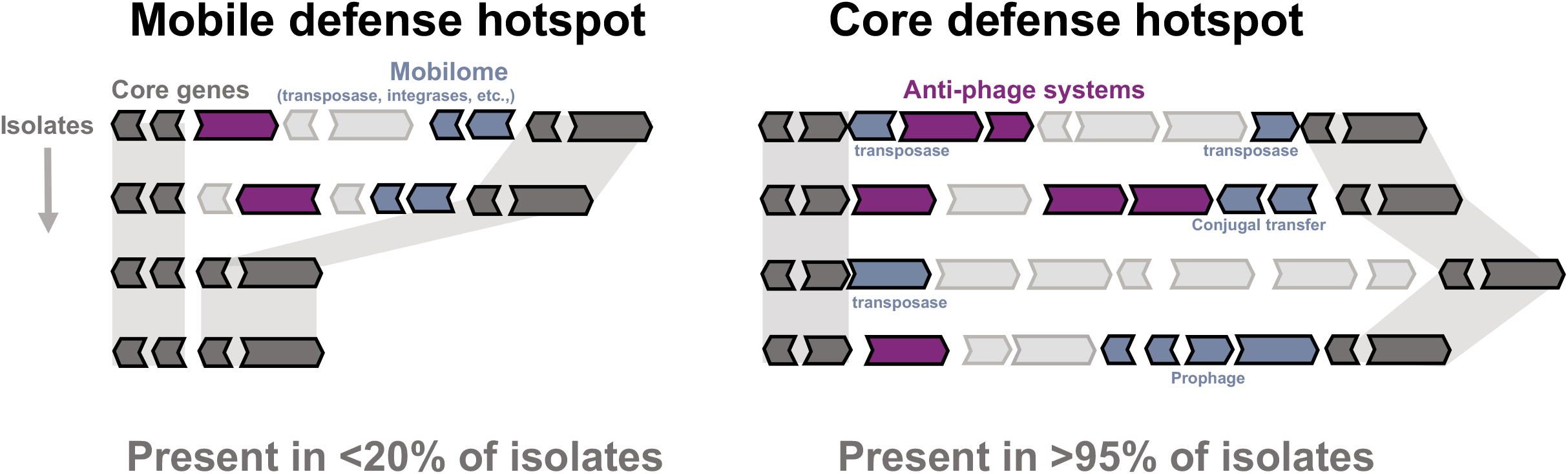

**Figure S1:**
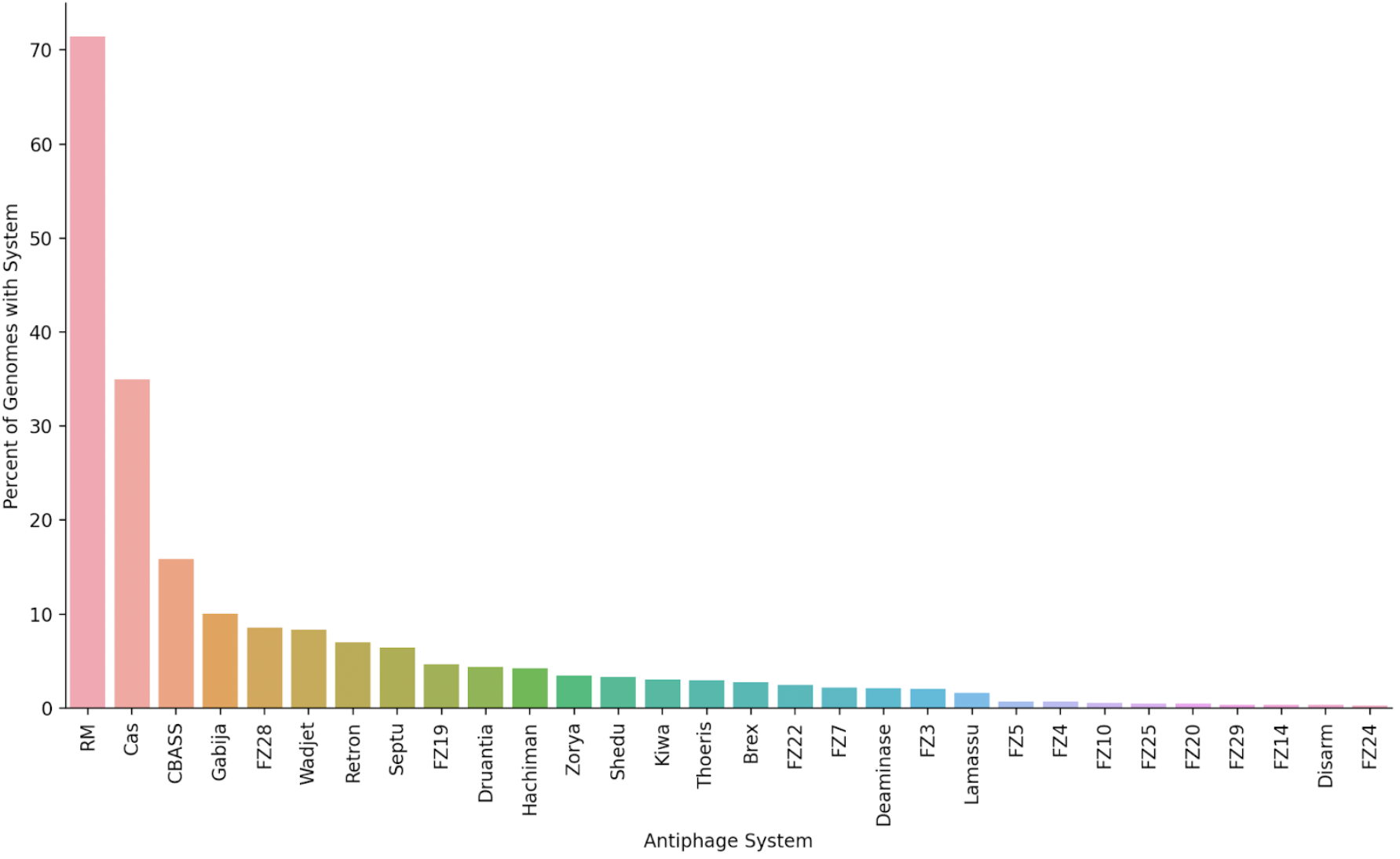
The abundance of each antiphage system in all genomes surveyed by ISLAND. This figure shows the fraction of all the genomes surveyed by ISLAND that harbor at least one representative of a given antiphage system.

**Figure S2:**
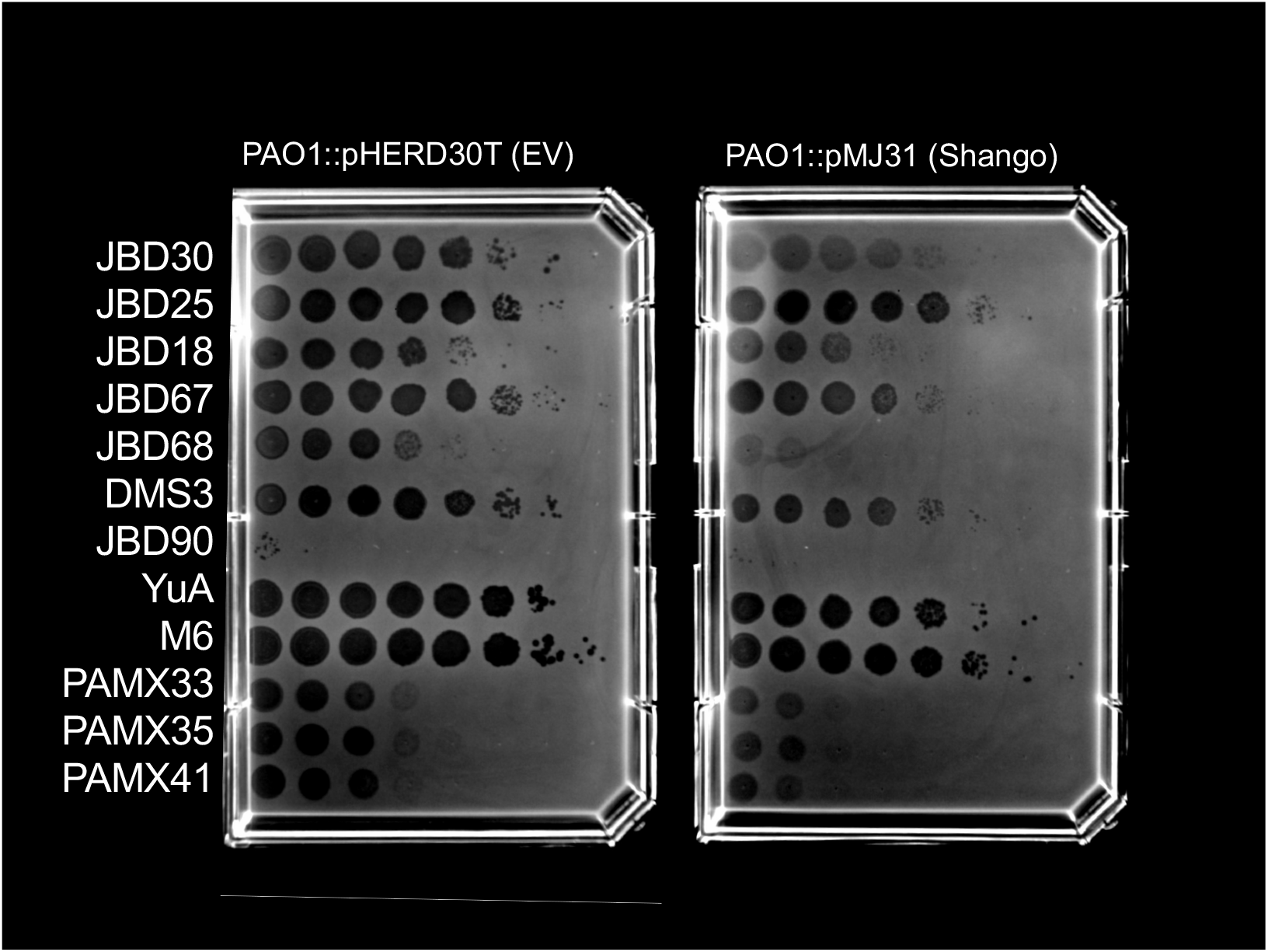
Shango phage assay. Several phage plated on PAO1::EV and PAO1::Shango strains.

**Figure S3:**
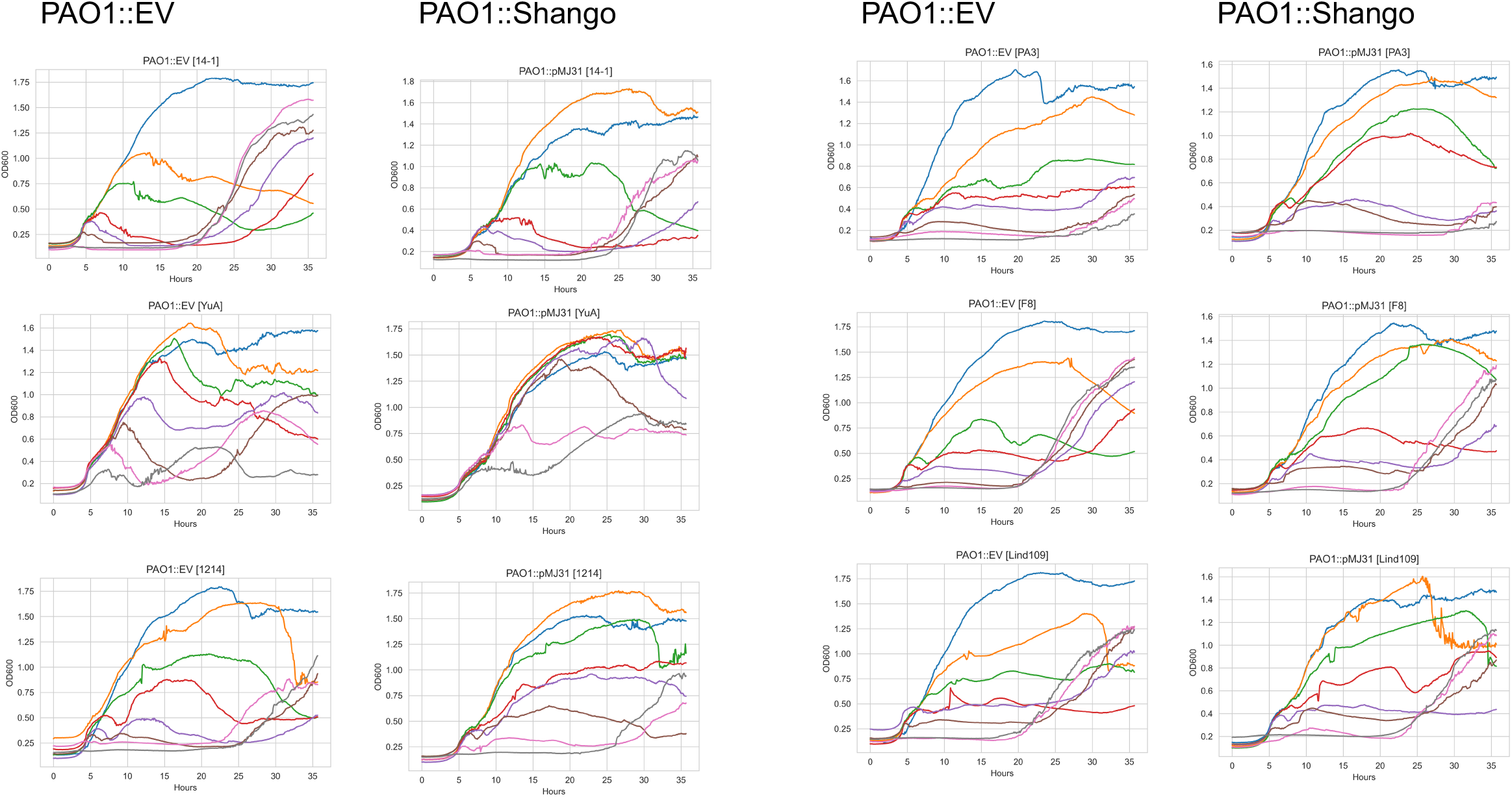
Growth curve assay. Liquid infection assay with PAO1::EV and PAO1::Shango infected with phage. Strains were grown in a plate reader with OD600 tracked over time.

**Figure S4:**
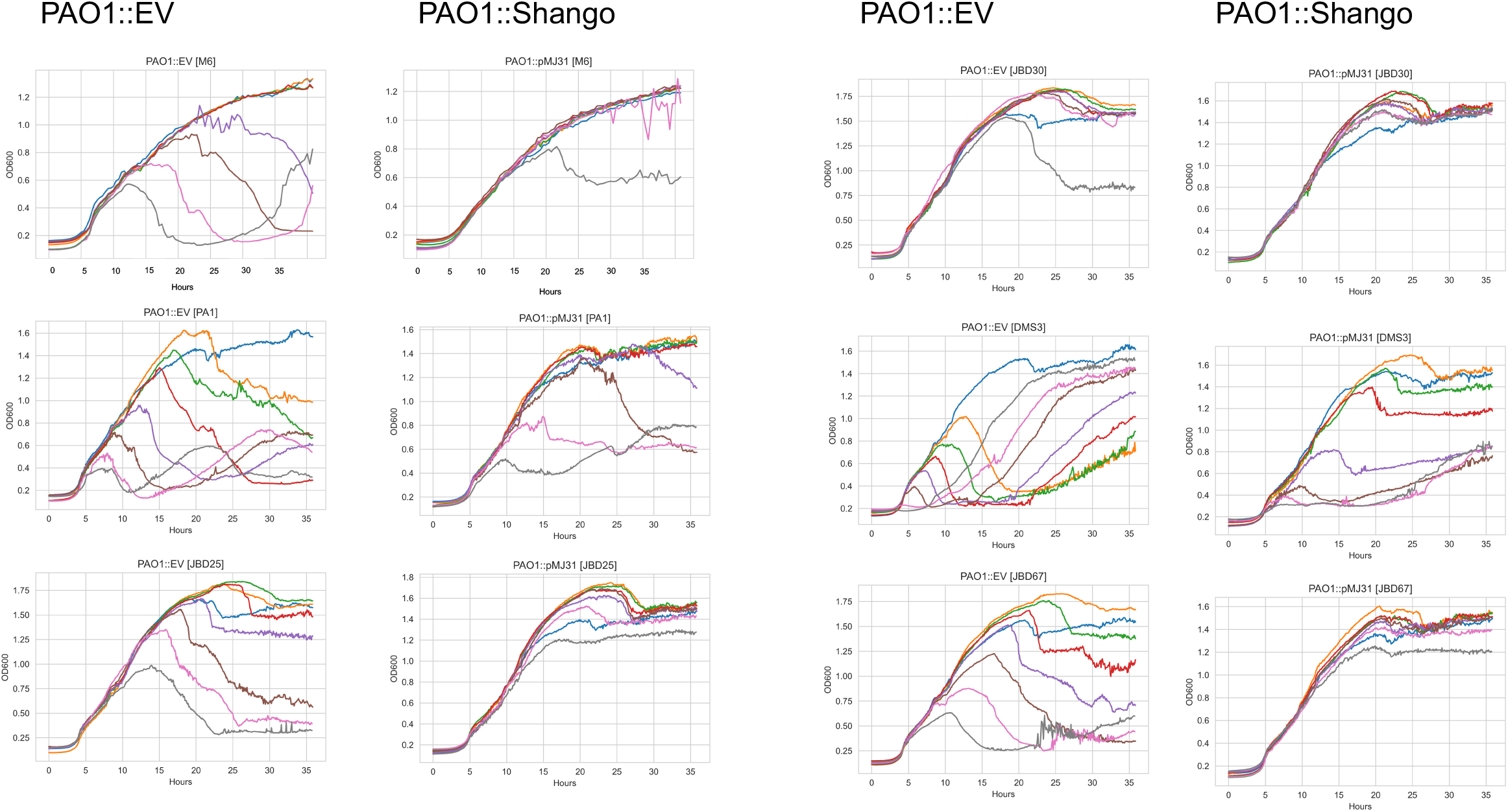
Growth curve assay (Cont.). Liquid infection assay with PAO1::EV and PAO1::Shango infected with phage. Strains were grown in a plate reader with OD600 tracked over time.

## Notes

https://doi.org/10.5281/zenodo.7254690

